# Computer Simulations of the Humoral Immune System Reveal How Imprinting Can Affect Responses to Influenza HA Stalk with Implications for the Design of Universal Vaccines

**DOI:** 10.1101/346726

**Authors:** Christopher S. Anderson, Mark Y. Sangster, Hongmei Yang, Sidhartha Chaudhury, David J. Topham

## Abstract

**Background:** Successful vaccination against the H1N1 Influenza A virus has required the continuous development of new vaccines that are antigenically similar to currently circulating strains. Vaccine strategies that can increase the cross-reactivity of the antibody response, especially to conserved regions, are essential to creating long-lasting immunity to H1N1 viruses. How pre-existing immunity affects vaccine-induced antibody cross-reactivity is still not well understood.

**Methods:** An immunological shape space of antigenic sites of hemagglutinin (HA) was constructed using viral sequence data. A Gillespie Algorithm-based model of the humoral immune system was used to simulate B cell responses to A/California/07/2009 (CA09) HA antigen after prior immunization with an antigenically similar or dissimilar strain. The effect of pre-existing memory B cells and antibody on the resulting antibody responses was interrogated.

**Results:** We found increased levels of highly-cross-reactive antibodies after immunization with antigenically dissimilar strains. This increase was dependent on pre-existing memory B cells. Furthermore, pre-existing antibody also interfered with the cross-reactive antibody response, but this effect occurred irrespective of the priming antigen.

**Conclusion:** These findings suggest that vaccination by divergent strains will boost highly-cross-reactive antibodies by selectively targeting memory B cells specific to conserved antigenic sites and by reducing the negative interference caused by pre-existing antibody.

## Introduction

Vaccination strategies are needed that induced highly-cross-reactive antibodies capable of binding a broad range of antigenically distinct influenza virus strains[1]. The seasonal H1N1 influenza virus vaccine requires continuous updating to compensate for antigenic drift in order to be effective[2]. Due to antigenic shift, the season H1N1 vaccine was not effective against the 2009 “Swine flu” pandemic[3]. In order to overcome these challenges novel vaccination strategies are needed[4].

The 2009 pandemic virus vaccine was capable of significantly boosting cross-reactivity, but only in younger age groups[5]. Serum antibody from these individuals bound a broad range of seasonal viruses and highly divergent avian influenza strains[5,6]. Additionally, there was an induction of antibodies towards the conserved stalk region of HA[6,7]. Additionally studies have suggested a role of Memory B cells in these highly cross-reactive antibody responses[8,9].

The age-specific differences seen after vaccination (or infection) with CA09 are thought to result from differences in the strains individuals have been previously exposed. Both pre-existing antibody and memory b cell specificity have been implicated in interfering with the cross-reactivity of secondary immune responses[9–14]. How pre-existing antibody and memory B cells contributed to the differences in cross-reactivity seen in different age groups is not well understood.

To gain insights into the role of memory B cells and pre-existing antibodies on cross-reactivity, we simulated secondary immune responses to CA09 HA antigen after priming with antigenically similar or distinct HA antigens. We investigated the effect pre-exposure to antigenically distinct antigens had on the cross-reactivity of a secondary antibody response.

## Materials and Methods

### HA Protein Sequences

Influenza HA protein sequences used in the model were obtained from Genbank: A/California/07/2009 (CA09) [NC_026433], A/Brisbane/59/2007 (BR07) [KP458398], A/South Carolina/01/1918 (SC18) [AF117241], A/Beijing/262/1995 (BE95) [AAP34323], A/Brazil/11/1978 (BR78) [A4GBX7], A/Chile/1/1983 (CH83) [A4GCH5], A/New Caledonia/20/99 (NC99) [AY289929], A/Singapore/6/1986 (SI86) [ABO38395], A/Solomon Islands/3/2006 (SI06) [ABU99109], A/USSR/90/1977 (US77) [P03453], A/New Jersey/11/1976 (NJ76) [ACU80014].

### Model Description

Simulations were performed using a Gillespie algorithm and a set of rate equations. The rate equations represent biological processes of B cells that occur during exposure to viral antigen. Together, these biological processes represent a simplified humoral immune system that can responds to antigen, produce antibodies, and remove antigen from the system. The model is identical to those described by Chaudhury et al. [15] accept for two modifications: (1) the number of antigenic sites representing each antigen was increased from 2 to 6 (2) long-lived plasma cells were added to the model [9] (Fig 1; Sup. Methods).

**Fig 1.**
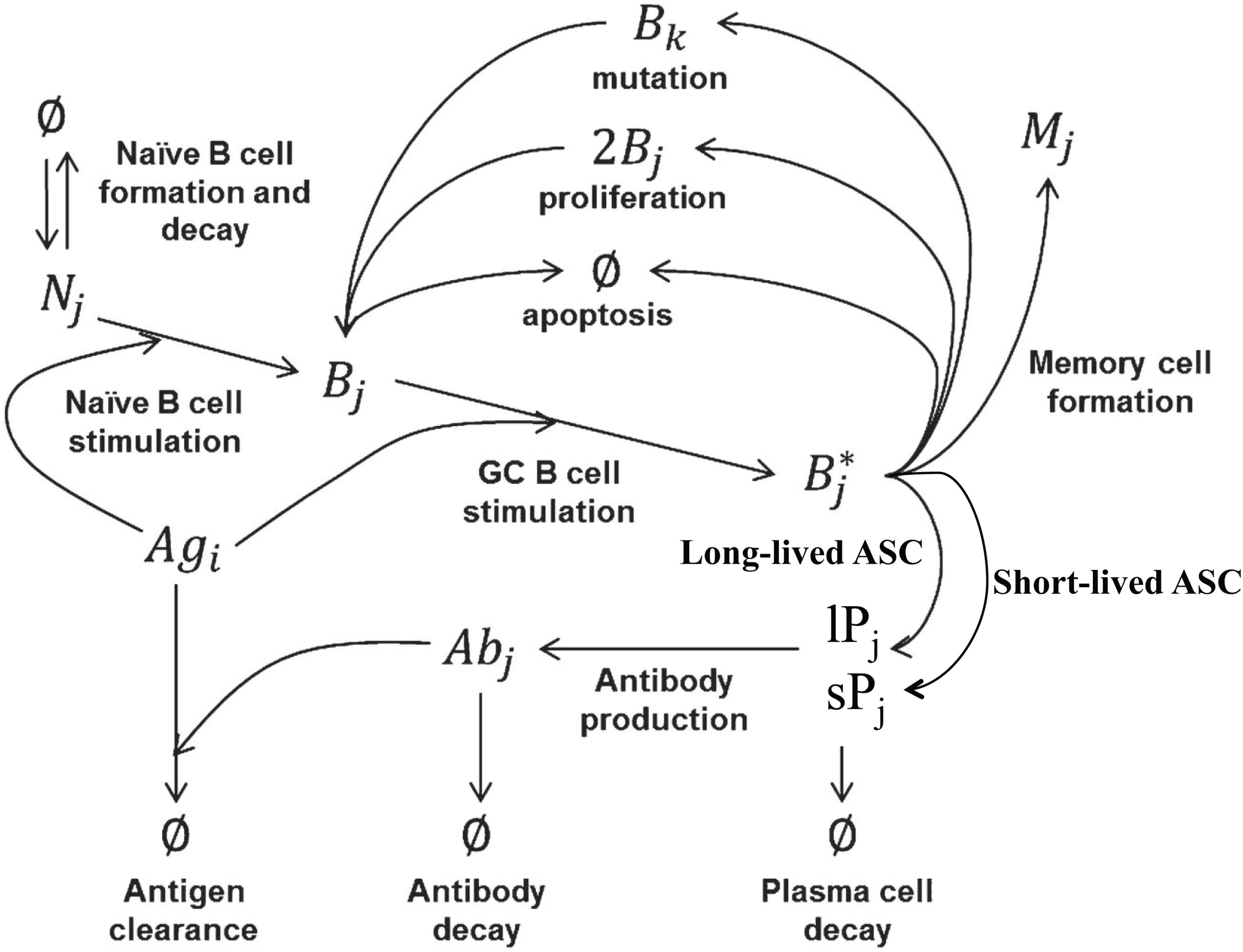
Schematic Representation of Humoral Immune System Model. Schematic is adapted from Chaudhury et al. [15]

### Simulating Immune Responses to the 2009 Pandemic Vaccine

Two scenarios were modeled with 50 simulations carried out for each scenario. In scenario one, the model was “primed” (antigen added to the system) with the SC18 HA antigen, then the immune response was allowed to resolve for 365 days, during which antibody level to returned close to baseline and subsequently immunized with CA09 HA antigen (Fig 2A). Scenario two was identical to scenario one except the model was primed with the BR07 HA antigen. B cell and antibody counts, genotype, and antigen specificities were tracked during the simulation allowing quantification of antigenic-site-specific B cells and antibodies during the simulation.

**Fig 2.**
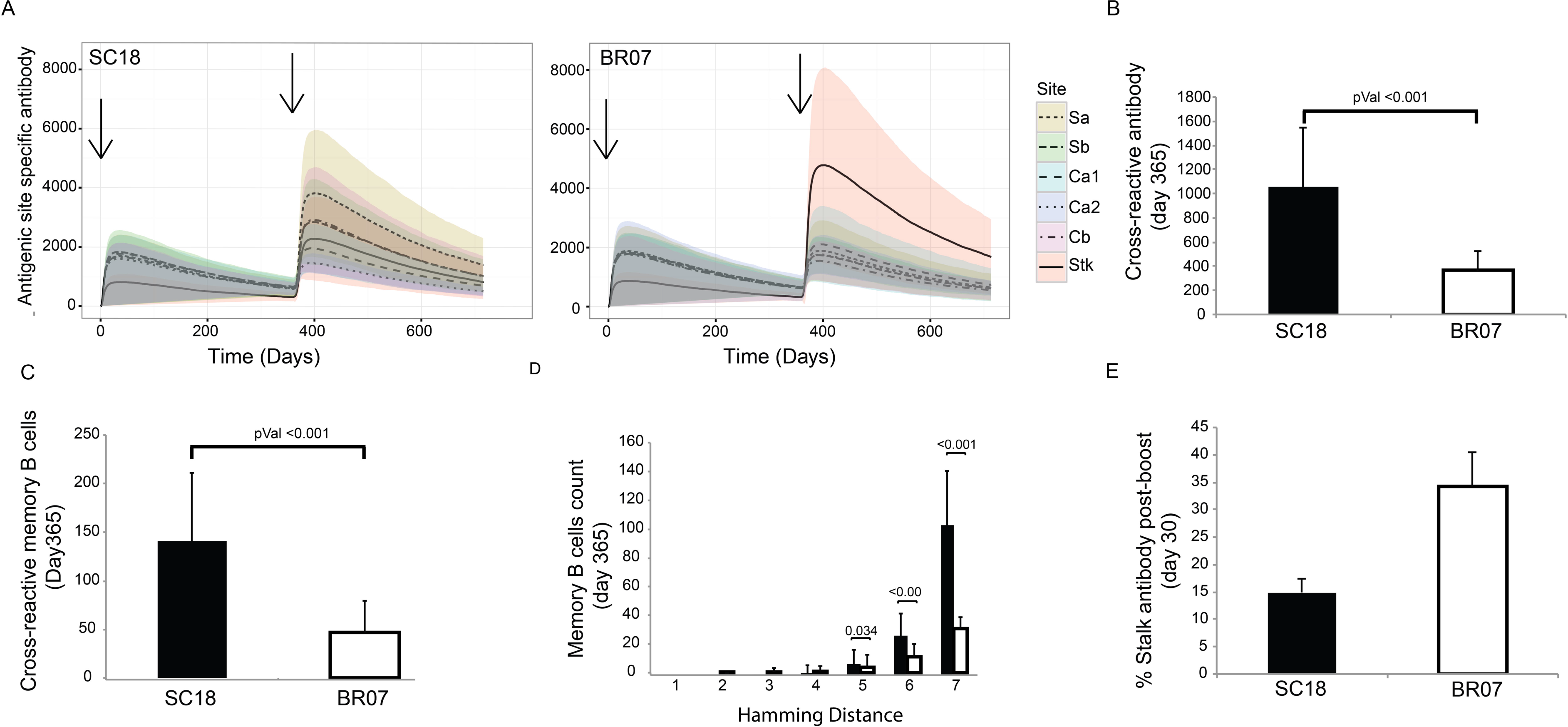
Immune Responses After Prime and Boost. (A) Antigenic-site-specific antibody titers to the priming antigen for the SC18 primed, CA09 boosted group (left) and BR07 primed, CA09 boosted group (right). Curves represent average titers for 50 simulations and colored area represents the standard deviation. Arrows represent times simulation was primed and boosted. (B) Average antibody titers and (C) memory B cells cross-reactive to CA09 pre-boost (Day 365). Statistic represents result of two sample t-test. (D) Affinity (antigenic distances 1-7) of memory B cells to CA09 HA antigen. Statistic represents result of two sample t-test. (E) Percent of stalk antigenic site-specific antibody for each priming group.

### Perturbed Models

Two perturbed models were created in order to assess the contribution of pre-existing antibody and memory B cells on the increase in stalk-specific antibodies. For the first model (“No Clearance”), the antibody clearance rate equation (which removes antigen form the system) was removed such that only basal decay of the antigen occurred. For the second model (“No Memory”), the memory B cell activation rate equation (which initiates germinal centers from memory B cells) was removed from the simulation such that only naïve B cells contributed to the germinal center reactions. Simulations were performed identical to the original model.

### 2009 H1N1 Vaccine Clinical Trial Human Serum

Samples from a previous clinical trial were used to test specific predictions of the simulations. The study was conducted under a protocol approved by the University of Rochester Research Subjects Review Board. Informed written consent was obtained from each participant. ClinicalTrials.gov identifier NCT01055184. Healthy adults and children were enrolled as previously described and results of this clinical trial have been published previously[6]. Subjects received a single intramuscular (i.m.) injection of inactivated influenza A/California/07/2009 (H1N1) monovalent subunit vaccine (Novartis). Each 0.5-ml dose contained 15μg of HA antigen. Administration of the vaccine (study day 0) took place from January 2010 to March 2010. Serum was collected before and 28 days after vaccination.

### Enzyme-linked Immunosorbent Assay

In order to test predictions of the model, we measured human serum antibody levels by Enzyme-linked immunosorbent assays (ELISA). Serum antibody levels were measured against recombinant HA proteins derived from historical antigens. ELISAs were performed using recombinant HA proteins coated on MaxiSorb 96-well plates (ThermoSci; 439454) overnight at 4°C. Plates were blocked with 3% bovine serum albumin (BSA) in phosphate buffered saline (PBS) for 1hr at room temperature. Serum was diluted 1:1000 in PBS/0.5% BSA/0.05% Tween-20. Plates were washed and incubated with alkaline phosphatase (AP)-conjugated secondary antibody for 2 hrs at room temperature. Plates were washed and developed using AP substrate (ThermoSci 34064). Recombinant HA proteins were obtained from Influenza Reagent Resource (Cat#: FR-67, FR-692, FR-65, FR-180, FR-699) and BEI Resources (Cat# NR-19240, NR-48873). Stalk specific antibody responses were measured using chimeric HA proteins that contained an “exotic” HA head but retained the conserved stalk region (cH9.1 and cH6.1). We report the relative change in antibody levels (d28/d0).

### Statistics

Two sample, two-tailed, t-test using the *t.test* function was performed using the base packages in R. A p-value of 0.05 or less was considered statistically significant. For group comparisons of the ELISA, a multivariate linear model (*lm* function in R) was performed to determine statistically significant differences between groups.

## Results

### Antigenic Distance Determination

We first determined the antigenic distance (AD) between HA antigens from 11 influenza virus strains. AD was determined using the H1N1 HA sequence-based antigenic distance approach previously described[16]. Given that each HA in the model contains 6 antigenic sites, and each antigenic site in the model contains 20 characters, the maximum epitopic distance (antigenic-site-specific antigenic distance; ED) is 20 and the maximum AD for each antigen is 120. 0verall, the real-life HA antigens of SC18 and CA09 had the greatest estimate of antigenic similarity with an AD of 21 (Table 1). BR07 and CA09 HA antigens had the greatest dissimilarity with an AD of 53. For SC18 and CA09 HA antigens, four of the five head epitopes had an ED of less than or equal to seven (Sa = 2, Sb = 3, Ca1 = 5, Ca2 =8, Cb = 3), with the Sa antigenic site having the least distance. Alternatively, BR07 and CA09 HA antigens had only one cross-reactive antigenic site (Sa = 8, Sb = 15, Ca1 = 7, Ca2 = 10, Cb = 13), with an ED of seven. Thus, in the model SC18 HA antigen was antigenically more similar to CA09 while the BR07 HA antigen was largely antigenically distinct from CA09.

**Table 1.**
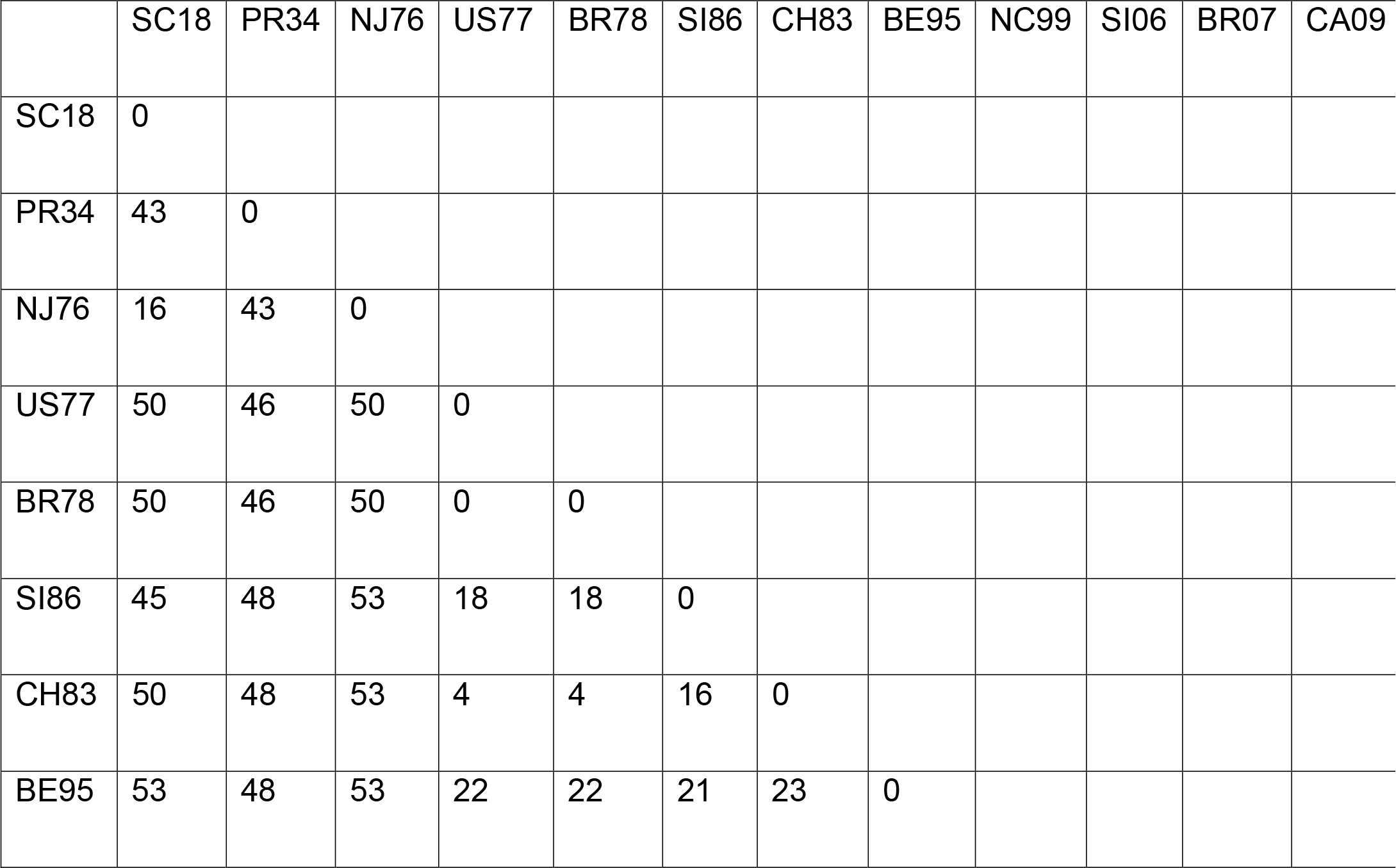
Antigenic Distances

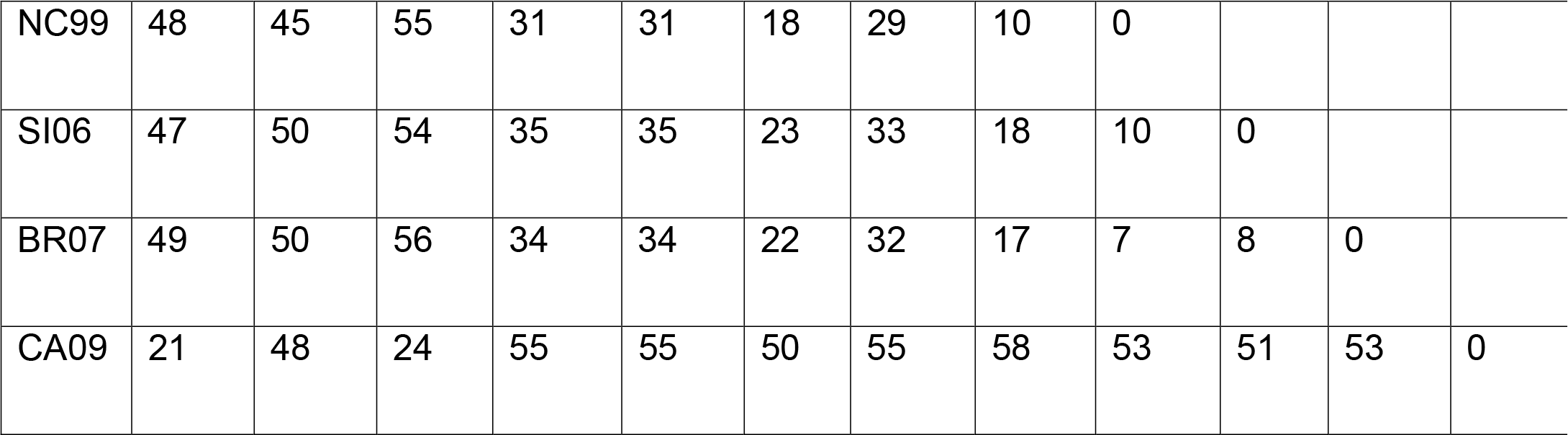

### Antigenic-Site-Specific Antibody Responses

Antigenic-site-specific antibodies and memory B cells reactive to the priming Ag (SC18 or BR07) were measured throughout the simulation. We found that counts of antibody and memory B cells specific for the priming antigen were similar across head antigenic sites and between groups (Fig 2A, S1 Fig A-D) and these similarities remained up until boosting (S1 Fig E-F). Stalk-specific antibody and memory B cell counts were significantly less than head antigenic sites making up about 10% of the total response and were similar for both groups (S1 Fig E and F). Therefore, antibody and memory B cell counts and specificities to their priming antigen were similar for both BR07-primed and SC18-primed groups.

Given that immune responses to the priming antigens were similar, we sought to determine if the cross-reactivity to CA09 differed between groups prior to boosting. We found a statistically significantly (p-values < 0.001) higher antibody and memory B cell levels cross-reactive to CA09 in the SC18 primed group compared to the BR07 primed group with a greater than 2-fold difference (Fig 2B-C). Although the amount of memory B cells with immunoglobulin receptors with low Hamming distance (high affinity) to CA09 HA antigen did not differ between groups, modest and high Hamming distance (low affinity) memory B cells were significantly increased in the SC18 group (Fig 2D). Taken together, priming-antigen-specific antibody and memory B cells were similar between groups, but were their cross-reactivity to CA09 was different.

After boosting with CA09 total antibody levels reactive to CA09 in the SC18 group were slightly higher, although this difference did not reach significance (S2 Fig A). For the SC18 primed group, antibodies to the Sa-antigenic-site of CA09 dominated with Sb and Cb antigenic-site specific antibodies also boosted (Fig 2A). Stalk antigenic site antibody were also boosted but to a lesser extent compared to head epitopes and comprised about 15% of the total antibody response (Fig 2E).

Unlike the SC18-primed group, stalk-specific antibody responses dominated for the BR07 primed group (Fig 2A) comprising 35% of the total antibody response (Fig 2E). A moderate increase in other antigenic-site-specific antibodies was also observed (Fig 2A). Antigenic site-specific differences between groups generally corresponded to differences in epitopic distances between the priming and CA09 antigens with those with closer epitopic distances showing a greater boost. Antibodies to the stalk antigenic site, which has the same epitopic distance in both groups, also demonstrated differences between groups. Despite being subdominant during priming, the large increase in stalk specific antibodies in the BR07-primed group demonstrates that the shorter ED at this site more than compensated for the decreased immunogenicity.

### Pre-Exposure Affects Cross-Reactivity of Secondary Responses

We found that after boosting with CA09 both groups had strong antibody responses to the antigens to which they had been previously exposed, but differed largely in responses to other strains (Fig 3A). This occurred despite the fact that antigenic distances to the 11 strains were not significantly different between BR07 and SC18 (two-sample t-test, p-value = 0.362). Generally, the SC18-primed group was cross-reactive to strains antigenically similar to CA09, while the BR07-primed group antibody response demonstrated greater cross-reactive to other strains. The BR07-primed group had a statistically significant increase in the number of stalk antibodies which were cross-reactive to all HA antigens (Fig 3B). Cross-reactive antibody levels in the SC18 primed group correlated well with the antigenic distance from CA09, while the BR07 primed group antibody cross-reactivity showed no linear correlation with antigenic distance (pval = 0.0001, pval = 0.4983; respectively). Therefore, although antigenic distance was a good predictor of cross-reactivity during the primary response of the simulation, for secondary immune responses antigenic distance alone was not sufficient to predict cross-reactive immune responses in individuals with different antigen exposure histories.

**Fig 3.**
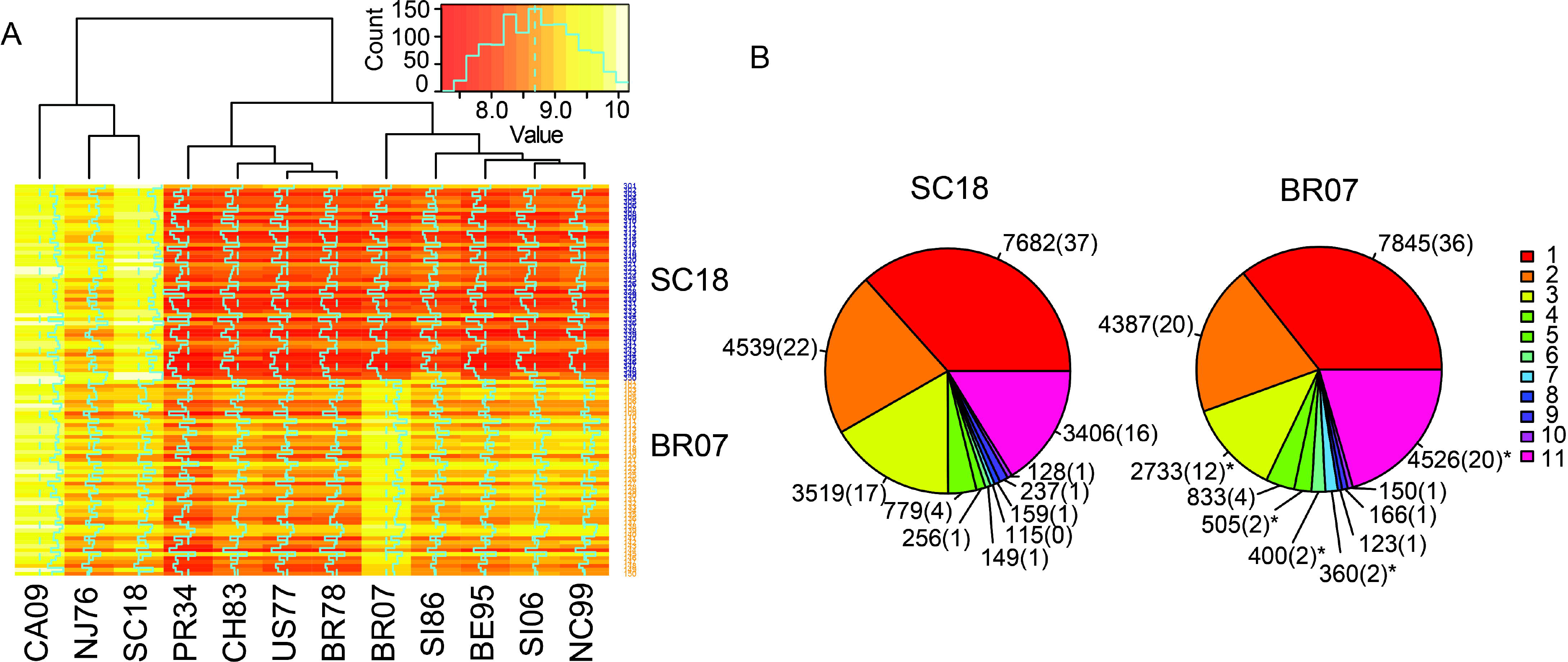
Cross-reactivity After Boosting with CA09. (A) Antibody levels to HA antigens from representative strains that circulated from 1918-2009 for the SC18-primed group or BR07-primed group. Antibody levels were taken at 30 days post-boost (day 395) and log transformed. Values are averages of 50 simulations. (B) Of all antibodies present at 30 days post-boost with CA09, pie-chart represents the number of those antibodies that are cross-reactive to other HA antigens (1-11 HA antigens). Number in parenthesis represents percentage rounded to the nearest whole number. Asterisk represents statistically significant difference (p-value < 0.05) between SC18 and BR07 groups as determined by two sample t-test.

### Contribution of Pre-Existing Memory B cells and Antibody on Cross-Reactivity

For the SC18-primed group, removal of memory B cell activation significantly increased antibody levels to the historical antigens (two sample t-test, p-value = 8.5 x 10^−26^) compared to the original unperturbed (“Normal”) model (Fig 4A). Interestingly, this increase occurred despite a decrease in stalk-specific antibody (Fig 4C). Removal of antibody clearance for the SC18-primed group also significantly increased antibody cross-reactivity (two sample t-test, p-value = 2.4 x 10^−15^), but this was to a lesser extent. For the BR07-primed group, removal of antibody clearance also significantly increased the cross-reactive response (two sample t-test, p-value = 4 x 10^−29^), but unlike the SC18-primed group, removal of memory B cells from the germinal centers statistically decreased (two sample t-test, p-value = 3 x 10^−76^) the cross-reactive response (Fig 4B). Stalk-specific antibody was also significantly increased in the “No Clearance” model, but significantly decreased in the “No Memory” model. Taken together, pre-existing antibody decreased cross-reactive antibody responses in both groups, while the effect of memory B cells had the opposite effect on cross-reactivity across groups. The increased cross-reactivity found in the BR07 primed group after boosting with CA09 resulted from activation of pre-existing memory B cells cross-reactive CA09 and not primarily from naive B cell responses.

**Fig. 4.**
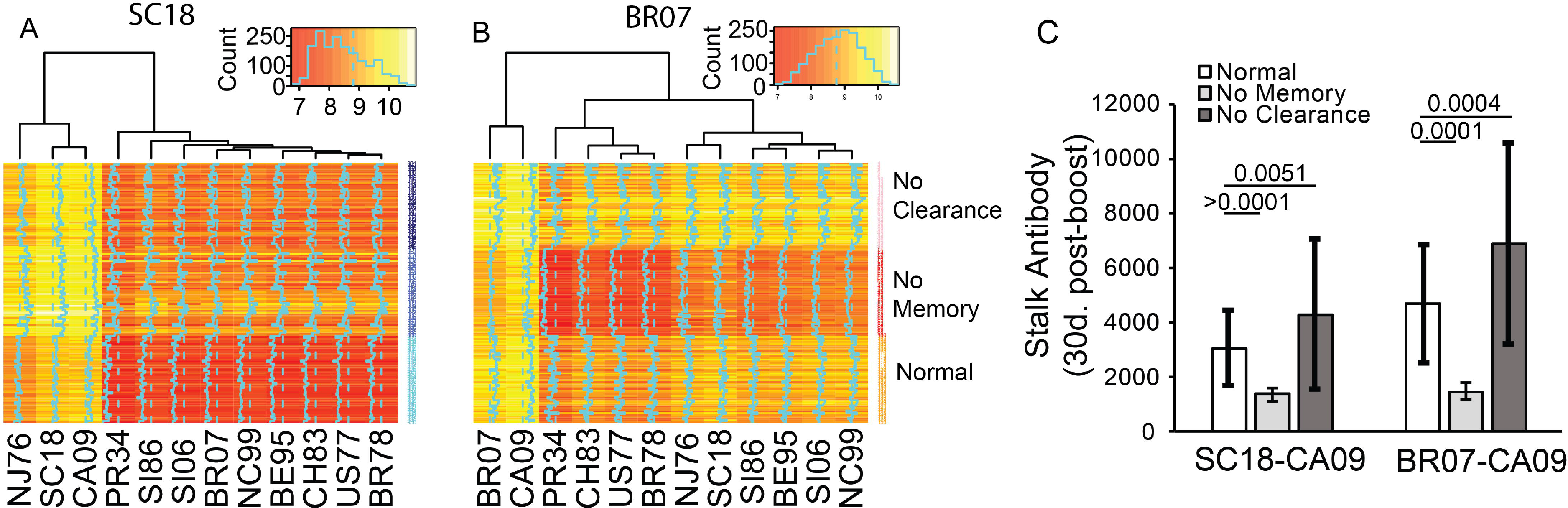
Antibody Levels After Perturbation of the Model. (A) Antibody levels measured at day 30 post-boost with CA09 for the SC18-primed group or (B) BR07-primed group for perturbed and normal models. Values were log transformed. (C) Stalk-specific antibody levels 30 days post boost with CA09 for each model. Average of 50 simulations, error bars represent standard deviation. Statistics were determined by two-sample t-test.

### Cross-Reactivity of Age-Stratified Serum

Finally, we sought to support our finding that cross-reactivity would be affected by prior exposure. Using serum from a previously described clinical trial, we measured antibody binding to a range of historical strains. Unsupervised clustering showed stratification by age group although this grouping was not exact (Fig 5A). Cross-reactivity antibody levels were generally increased in the 18-32-year-old age group, but not to all HA antigens (Fig 5B). Importantly, the HA antigen, CH6.1, showed a statistically significant increase in antibody levels for the 18-32-year-old age group compared to the 60+ age group (multivariate regression model, p-value = 0.04358). Alternatively, NC99 antibody binding was modestly increased in the 60+ age group, although the difference did not pass our significance threshold (multivariate regression model, p-value = 0.09916). Taken together, these finding support that cross-reactivity of secondary immune responses are affected by exposure history.

**Fig 5.**
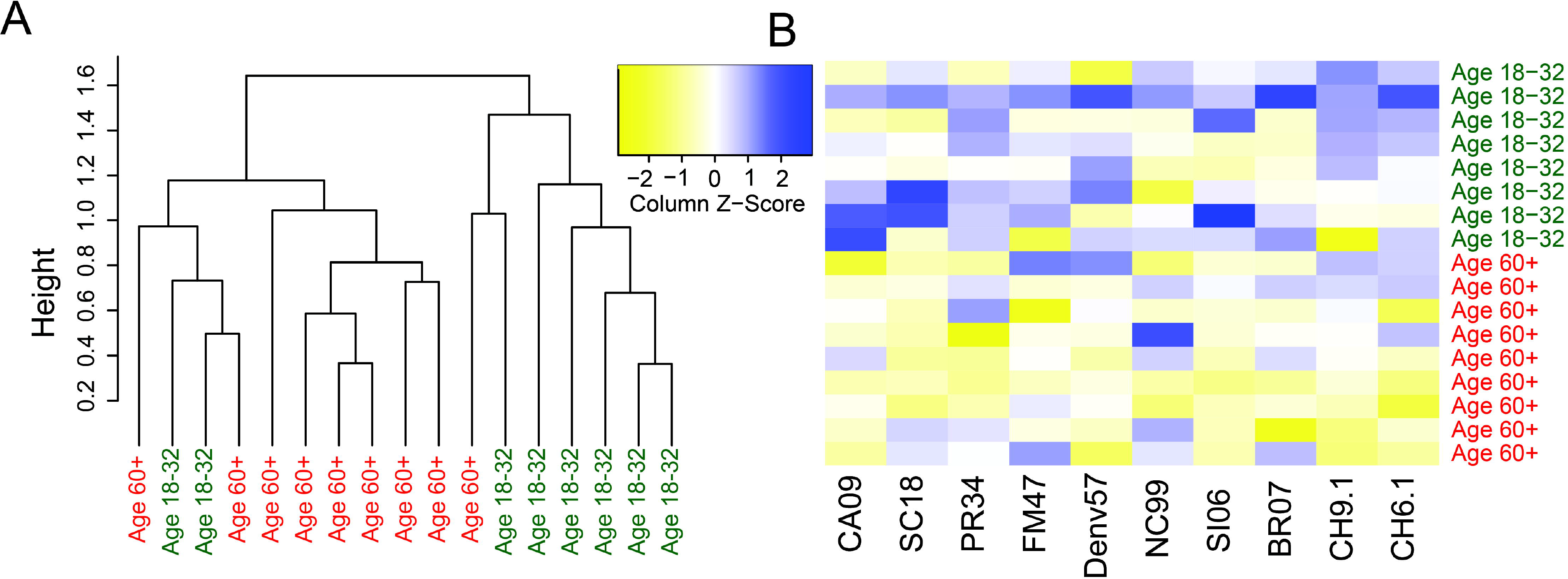
Serum Antibody Levels for Age-Stratified Cohort. (A) Hierarchical clustering of antibody binding measured against recombinant HA proteins using ELISA. (B) Heatmap of ELISA antibody binding data. Data represents the relative change in binding. Data was log transformed and standardized, values represent column z-scores

## Discussion

Protection against antigenically drifting or shifting H1N1 viruses by vaccination requires continuous reformulation of the vaccine. Increasing the cross-reactivity induced by vaccination is a hallmark of “universal” vaccine efforts[4]. Here we computationally simulated immunization with SC18 HA antigen or BR07 HA antigen and subsequent immunization with CA09 HA antigen. We also evaluated the effect of pre-existing antibody and memory B cells on the levels and cross-reactivity of antibodies after immunization with CA09 HA antigen.

Elucidating the combined effect of differences in HA antigenic site conservation, pre-existing immunity, epitope dominance, and B cell and antibody specificities is a daunting task for the experimentalist. However, computational models allow explicit manipulation and observability of biological processes that are not possible with typical animal and human models. Perelson et al. hypothesized that B cell receptor repertoires (paratopes) exist in an immunological shape space and antigen binding differences between them are represented as distance in shape space[17]. Smith et al. subsequently derived the parameters of such an immunological shape space for influenza viruses[18]. Moreover, Smith et al. developed a computational model of the humoral immune system and demonstrated that such a model can be used to understand secondary immune responses to influenza antigen[9]. Recently, Chaudhury et al. developed a stochastic simulation model using the parameters developed by Smith et al. and expanded the model to include multiple antigenic sites of different conservation[15]. Here we adapted the Chaudhury et al. model to better simulate immune responses to the H1N1 influenza HA antigen.

In the current study, we aimed to understand how prior exposure to influenza virus antigens affects the antibody specificity during secondary exposures. This work was an extension of the work originally performed by Smith et al.[19] and the theory of Shape Space originally developed by Perelson et al.[17]. Consist with Smith et al. findings, we found that the antigenic relationship between the first and secondary exposure antigens largely affect the specificity of the antibody response. Moreover, during secondary immune responses in the model, antigen was removed from the system more quickly in the group previously exposed to an antigenically similar strain during the primary exposure, consistent with the notion of antibody mediated negative interference[9]. Moreover, the increase in stalk-specific antibodies in the BR07 group during boosting is consistent with weakening of negative interference from head-specific antibodies. Additionally, the increased antibody response to the CA09 strain in the SC18 exposed group after boosting supports the notion of positive interference, in which antibody responses from pre-existing memory B cells are increased. Taken together, our findings are consistent with the Antigenic Distance Hypothesis described by Smith et al.[9].

The expansion of the Shape Space-based model to include multiple antigenic sites by Chaudhury et al. was a major advancement in use of the model to understand B cell specificity across complex antigens[20]. By incorporating multiple antigenic sites, the model creates competition for antigen between B cells complementary to different antigenic sites on the same antigen. Although Chaudhury et al. modeled a multivalent vaccine, our findings are consistent with their finding that antibody responses to a normally sub-dominant stalk antigenic site will dominate when the antigenic distance between head antigens are large, even when immunogens are given successively. Additionally, the large increase in stalk-specific antibodies in the BR07 group is consistent with reports on universal vaccine development that apply a similar strategy to boost stalk specific antibodies[4,21].

One of the most significant findings of the 2009 pandemic was the ability of 2009 pandemic vaccine to induce antibodies able to bind antigenically distinct viruses[22,23]. The results of our simulations corroborate these findings by demonstrating that BR07-primed individuals had increased antibody reactivity to 1918-like HA antigens (CA09, SC18, NJ76) as well as seasonal H1N1 viruses after exposure to CA09, while SC18-primed individuals produced antibodies primarily reactive to 1918-like HA antigens. These findings are consistent with the reports suggesting that original virus exposures, not age, affected the vaccine response to the 2009 vaccine[5,24]. Furthermore, although only slightly different, SC18 antibody titers were higher than CA09 titers after boosting with CA09 in the SC18 group but CA09 titers were higher than BR07 in the BR07 primed group. The SC18-primed group demonstrated antibody responses consistent with original antigenic sin (OAS), where individuals display antibody levels highest against viruses that circulated when they were young (and first exposed to influenza) despite subsequent exposure to drifted influenzas viruses[25]. The BR07-primed group did not show the same OAS phenotype as CA09 antibodies were highest after boosting, but antibody levels to BR07 were highest in the BR07 exposed group.

Pre-boost antibody levels of the SC18 primed group were almost 3-fold greater for CA09 than those primed for BR07, similar to what has been reported[26]. Additionally, the fold-change increases in the antibody response to the stalk is consistent with published reports[6]. We found that the antigenic site (Sa), which had the least antigenic difference among SC18 and CA09 HA head antigenic sites, dominated the antibody response after boosting with CA09 in the SC18-primed group. The Sa antigenic site dominance is the SC18 group is consistent with experimental data showing that antibody responses from the 60+ year old individuals had antibody responses focused on the Sa site of CA09[27]. Furthermore fold change titers (pre-boost/post-boost) were decreased in the SC18 primed group suggesting it is important to take into account priming history of the elderly when trying to assess immunosenescence or predict responses in different age groups[9,11,28,29].

Here we showed that cross-reactivity can be boosted with sequential immunization by antigenically distinct antigens. Differences in cross-reactivity after sequential vaccination have been previous demonstrated in the context of pandemic virus vaccines where more highly cross-reactive antibodies were observed in subjects primed with an A/Hong Kong/97 H5 vaccine and later boosted with an A/Vietnam/04 vaccine, who then subsequently mounted antibody responses recognizing both vaccine strains, as well as a third H5 strain (A/Indonesia/05) not included in either vaccination [30].

It is worth noting that we found a decrease in stalk specific antibodies when the number of head antigenic sites was increased. This finding may help answer how the ratio of head to stalk epitopes of HA affects the subdominance of the stalk antigenic site. If in reality the head contains more antigenic sites than the stalk, the model predicts that stalk-antigenic site response will be decreased. Our analysis suggests that stalk specific antibody truly decreases with the addition of head antigenic sites, and it was not that stalk-specific antibodies remain constant and only the relative amount compared to the head is changed. It also suggests that the immunologic subdominance of the stalk does not necessarily mean it is inherently less immunogenic, having implications for targeting this domain in universal vaccination efforts.

Lastly, the work described here demonstrates the limitations with the current vaccine selection process that relies only on antigenic and phylogenetic distances between strains. Here, the shorter antigenic distance between SC18 and CA09 compared to BR07 and CA09 led to two different outcomes. For instance, the SC18 primed group had low titers to US77 after boost with CA09, while the BR07 primed group had greater titers. Therefore, although the antigenic distance between CA09 and US77 is fixed, previously exposed individuals produce antibody responses inconsistent with these antigenic distance estimates. This suggests that serum samples are not ‘impartial observers’ of antigenic similarity and they are highly biased by their own immune histories. This is an inherent challenge with the current influenza vaccine approach and highlights the need to consider prior exposure histories when trying to predict antibody specificities after vaccination.

In conclusion, our findings are consistent with other studies that point to negative and positive interference as a mechanism affecting this enhancement of cross-reactivity after sequential immunization[9–11,31]. Therefore, immunization regimens that can relieve negative interference while increasing positive interference (especially to conserved regions on an antigen) may act to broaden cross-reactive immunity to the Influenza virus[15,32].

## Acknowledgments

We thank the Center for Integrated Research Computing and the Health Sciences Center for Computational Innovation for computational assistance and resources. Thank you Carrie A. Anderson and Elaine Smolock for help with the manuscript. Thank you Alan Perelson, Martin Zand, Juilee, Thakar, John Treanor, Jim Miller, and Paige Lawrence for supervision of this work. Thank you Derek Smith for helpful comments early on in this project. Thank you Anthony DiPiazza, Matt Brewer, and Patrick McCall for discussions on the project. Chimera proteins were a kind gift from Dr. Florian Krammer. Funding for this work was supported by the New York Influenza Center of Excellence NIH/NIAID/DMID, HHSN272201400005C and the University of Rochester Pulmonary training grant T32-HL066988.

---

**Supplimental Figure 1:**
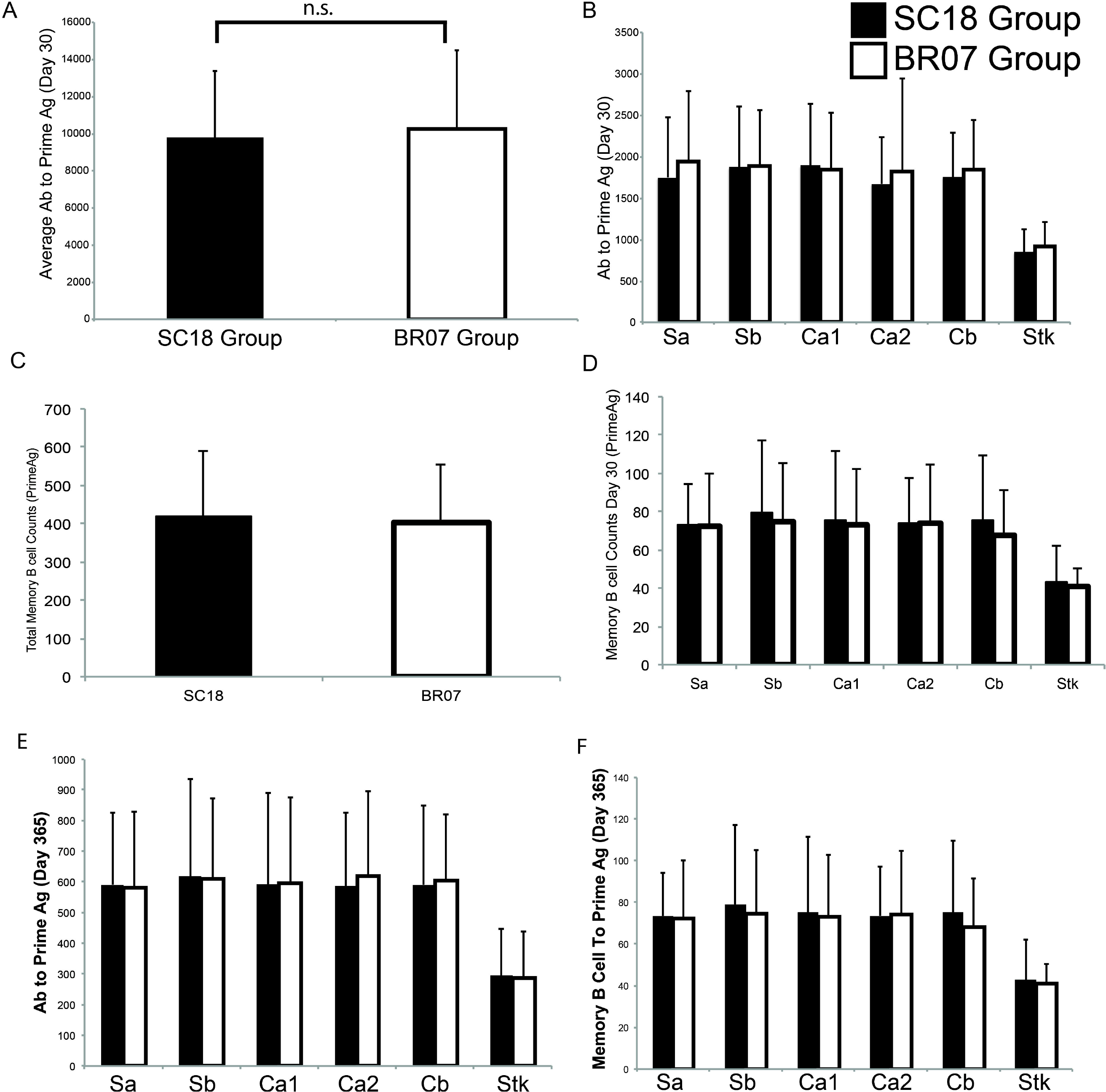
Immune Responses To Priming Antigen. (A)Representative simultion of B cell, antibody, and antigen kinetics after priming for SC18 primed group (B)or BRO? primed group. (C)Average antibody titers to priming antigen measured at day 30 post immunization tion for each group. Two groups of 50 simulations were immunized with either (SC18 or BRO?). (D) Epitope-specific antibody titers to priming antigen for each epitope in the model, day 30. (E) Average counts of memory B cells specific to the priming antigen for each group measured at day 30. (F) Epitope-specific memory B cell counts specific to prirming antigen measured at day 30.(G) Epitode-specific anti-body titers to priming antigen and (H) memory B cell measured pre-boost (Day 365)

**Supplimental Figure 2.**
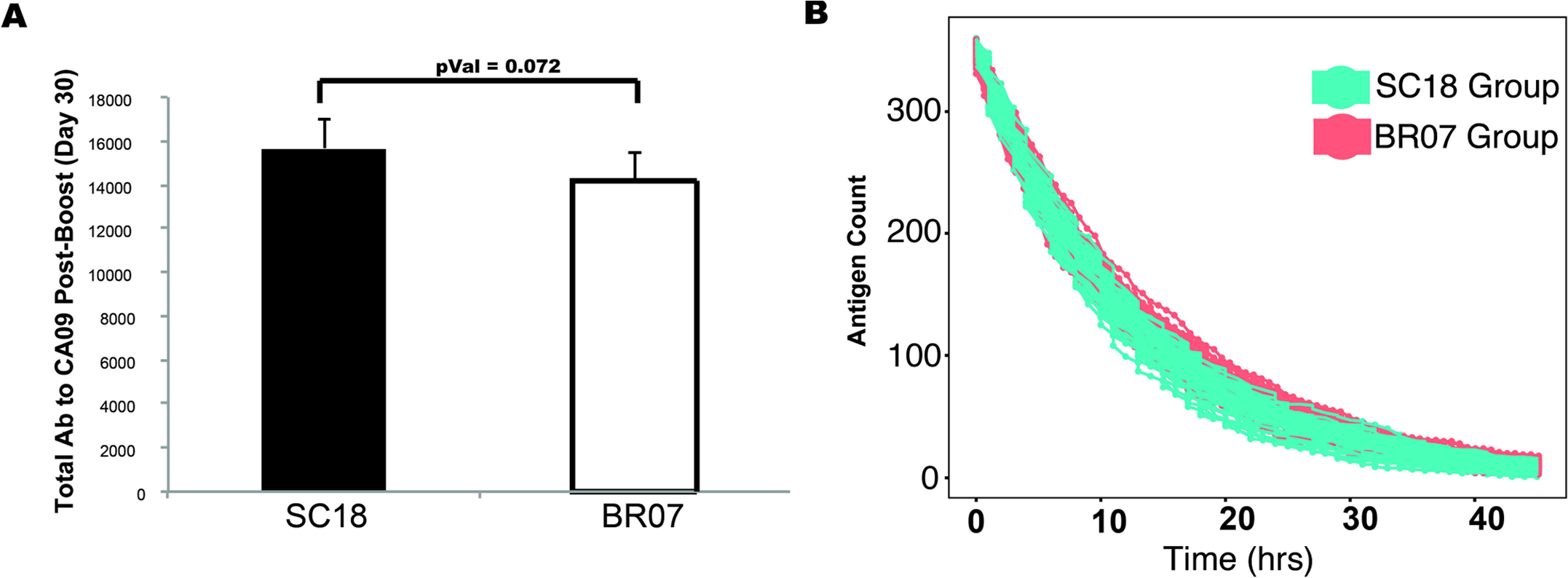
(A) Total antibody to CA09 measured day 30 after boosting with CA09. (B) Antigen clearance kinetics post-boosting with CA09 antigen for all subjects in each priming group.

