## Supplemental Methods for "Computer Simulations of the Humoral Immune System Reveal How Imprinting Can Affect Responses to Influenza HA Stalk with Implications for the Design of Universal Vaccines"

### Representing HA Antigen in the Model

In order to model the properties of antigen-immunoglobulin binding, we used the principals of Immunological Shape Space originally theorized by Perelson et al.[1]. Parameters values of the Shape Space Model were set to those previously determined by Smith et al. [2] and Chaudhury et al [3].

The H1N1 Hemagglutinin (HA) antigens were modeled using the following criteria: Each HA antigen contained 5 distinct, equally dominant, antigenic sites representing the 5-canonical head antigenic sites of H1N1[4]. We acknowledge that equal immunodominance among the 5-canonical head antigenic sites may not be exact, but were modeled this way for simplicity. In addition to the 5 head antigenic sites, each HA contains a single, subdominant, stalk antigenic site[5]. Each antigenic site is symbolically represented in the model using 20-character strings made up of 4 unique symbols (e.g. “AAAAABBBBBCCCCCDDDDD”).

Antigenic differences across antigenic sites of HA antigens in the model was determined using a virus sequence data using a method similar to previously described method [6]. This approach uses virus sequence data to estimate the epitopic distance (antigenic-site specific antigenic distance) between regions of HA[6]. In brief, for each virus strain used in the model, the HA protein coding regions of the RNA sequence data was virtually translated into protein sequences. Then, for each head antigenic site (Sa, Sb, Ca1, Ca2, Cb) protein sequences are truncated to include only the amino acids comprising that antigenic site. The Hamming distance (i.e. number of amino acids differing between the truncated sequences) is then calculated between all antigenic sites across all strains. To normalize for differences in the number of amino acids comprising each antigenic site, the Hamming distance was divided by the total number of amino acids comprising each antigenic site. The resulting value was then multiplied by 20. The result is that each antigenic site has a distance (range 0 to 20) to ever other antigenic site for every strain used in the model. These antigenic site-specific distances were realized in the model by randomly generating 20-character strings until the Hamming distances between virtual HA antigenic sites matched the distance calculated from the viral sequence data above. Therefore, each HA antigen in the model is represented as six (5 head, 1 stalk) independent 20-character strings (e.g. CA09 Sa antigenic site = “BBAAABBBBBCCCCCDDDDD”).

Unlike head antigenic sites, the exact number and location of the HA stalk region all possible antigenic sites are still largely unknown. Studies have demonstrated there are at least 1-2 epitopes in this region[5], with antibodies directed to the fusion domain having the ability to affect infectivity of the virus. We therefore chose to model a single HA stalk antigenic site (Stk). The stalk region of HA is more highly conserved amongst H1N1 viruses compared to head epitopes. Therefore, for simplicity, we chose to model the stalk antigenic site as a completely conserved antigenic site, although it is likely that multiple antigenic sites exist in the stalk region and vary in conservation[7,8].

Immunogenicity parameters for HA antigenic sites were chosen from experimental mouse studies[9,10]. These studies suggest that stalk-specific antibody responses make up about 20% of the total response[9,11]. Therefore, an immunogenicity parameter value for each antigenic site was used to account for these differences (Table 1). An immunogenicity parameter of one represents equal immunogenicity. Because the HA head is immunodominant compared to the stalk, immunogenicity parameters were adjusted to 0.8 for the stalk antigenic site and 1.2 for all head antigenic sites which resulted in a primary antibody responses in the simulation that was five-fold lower to the stalk antigenic site compared to a single head antigenic site, comparable to experimentally determined responses[9,11].

### Simulating the Humoral Immune System

Borrowing from work by Smith and Chaudhury, we chose to represent the immune response using a simplified version of the humoral immune system representing only the B cell arm. The model is comprised of agents of the immune system each grossly reflecting the higher-level functions of the humoral immune system. The model consists of 7 agents: Naïve B cells, stimulated B cells, germinal center B cells, short-lived plasma cells, long-lived plasma cells, memory B cells, and antibody (Fig 1).

Here we use the simulation method developed by Chaudhury *et al*. 2013[3]. This method uses a stochastic chemical-kinetics based approach[12,13] to simulate the progression of an immune response at a repertoire-scale. Components such as B cells, antigens, and antibodies are modeled as chemical species and biological processes such as antigen-binding, somatic mutations, or B cell replication are modeled as chemical reactions. All parameters in the model were set as previously described[3] with the exception of long-lived plasma cell parameters which were modeled as previously described[14](Table 1).

Immunoglobulin receptors for B cells are represented in the model by randomly generated, 20-character strings (e.g. “AABBCCDDAABBCCDDAAAA”) which reflects the stochastic nature of the B cell repertoire *in vivo*. The length and number of unique symbols at each locations gives the following properties: a potential immunoglobulin repertoire of 10^12^ B cells, a 1 in 10^5^ chance of B cells responding to a particular antigen, and an expressed repertoire of 10^7^ B cells [15-19].The affinity of each B cell for a given antigenic site is determined from the Hamming distance between the immunoglobulin string and the antigenic site string.

The simulation begins with the Naïve B cells at steady-state with equal formation and decay. When antigen is added to the model, Naïve B cells bind antigen and become stimulated B cells. Stimulated B cells form germinal centers and become germinal center B cells capable of stochastically differentiating into plasma cells (long or short lived) or memory B cells. Plasma cells secrete antibody capable of binding and removing antigen from the system. Memory B cells can also be stimulated by antigen and form germinal centers. Follicular helper T cells and antigen presenting cells are modeled implicitly.

**Table 1. Model Parameters**

| **Parameter** | **Symbol** | **Value** |
| --- | --- | --- |
| No. of naïve B cells |  | 5 x 10^7^ cells |
| Ag dose |  | 360 units |
| GC carrying capacity | 𝒌 | 5000 cells |
| Ag parameters |  |  |
| **Epitopes** |  | 6 |
| **Epitopes1** |  |  |
| Immunogenicity | 𝐲 | 0.8 |
| Clearance | 𝒑 | 1.0 x 10^-4^ cells |
| Epitopic distance |  | 0 |
| **Epitope 2-6** |  |  |
| Immunogenicity | 𝐲 | 1.2 |
| Clearance | 𝒑 | 1.0 x 10^-4^ cells |
| Epitopic distance |  | Varied |
| Intrinsic decay | gAg | (12 h)^-1^ |
| **B cell parameters** |  |  |
| B cell enhancement factor | 𝛆B | 10 |
| Ab enhancement factor | 𝛆Ab | 2.5 |
| Naïve B cell formation rate | 𝒌N | (4.6 x 105 h)^-1^ |
| Naïve B cell stimulation rate | 𝝈N | (1 d)^-1^ |
| GC B cell stimulation rate (base) | 𝝈base | (8 h)^-1^ |
| GC B cell stimulation rate (maximum) | 𝝈max | (15 min)^-1^ |
| GC B cell replication rate | r | (8 h)^-1^ |
| Mutation probability | μ | 0.1 |
| Differentiation probability | 𝜹 | 0.1 |
| Memory cell stimulation | 𝝈Max | (1 d)^-1^ |
| Ab production rate | 𝒌Ab | 1 |
| Naïve B cell decay rate | gB | (4.5 d)^-1^ |
| GC b cell decay rate (base) | gB | (4.5 d)^-1^ |
| Plasma cell decay rate | gP | (3 d)^-1^ |
| Ab decay rate | gAb | (10 d)^-1^ |

### Simulating Biological Processes

Simulation of biological processes defining the humoral immune system were carried out by describing a set of rate equations that describe the underlying biological reactions and then applying the Gillespie algorithm, a dynamic Monte Carlo method [3,20].

In the simulation, the binding affinity between the paratope and antigenic sites is proportional to the number of symbols that are complementary between them (their Hamming distance). Two antigenic sites cease to be cross-reactive when a 35% change or more in amino acid sequence in the amino acids that make up each HA antigenic site[21,22]. Therefore, there are eight degrees of cross-reactivity between epitopes corresponding to epitopic distances 0 through 7. Although epitopic distance ranged from 0 to 7, affinity ranged for each site can vary from 4 to 7 reflecting the degeneracy in paratope sequences that can achieve maximum binding affinity to their epitope. This range represents a 10^4^-fold difference in binding affinity of the immunoglobulin between naïve and fully matured B cells.

((1))	$Q_{\mathrm{ij}}=\{ \begin{matrix} 1, \\ \varepsilon^{\alpha-d\left( i,j \right)}, \\ 0, \end{matrix} \begin{matrix} d\left( i,j \right)< \alpha\\ \alpha\leq d\left( i,j \right)\leq\lambda\\ d\left( i,j \right)> \lambda\end{matrix}$

The binding affinity (Qij) between paratope i and the epitope j is a function of their Hamming distances, d(i,j) (eq. 1). Parameter ε is an enhancement factor that reflects the fold increase in apparent binding of B cells compared to antibodies (10 and 2.5, respectively). The increased avidity of B cells in comparison to a single antibody in the model reflects the many immunoglobulin molecules found on the surface of B cells resulting in many possible interactions between B cells and antigen[3]. Parameters, *α* and *λ*, were kept consistent with the Smith et al. and Chaudhary et al. models and represent the minimum and maximum Hamming distance (4 and 7, respectively), but it should be acknowledged other methods exist that distinguish Hamming Distances below 3[23].

((2a))	$N_{j} \overset{\to}{P_{Ag}gN} 0$

((2b))	$0 \overset{\to}{k_{N}} N_{j} k_{N}=5 \times{10}^{7}\cdot P_{Ag}gN$

The model simulates an animal size B cell repertoire of 10^7^-10^8^ B cells [18,19]. The life expectancy of unstimulated (naïve) B cells is 4.5 days. Naive B cells with randomly generated immunoglobulin (character-strings) are created such that there is steady population of 5 x 10^7^ naive B cells per antigen and the numbers of naïve B cells are balanced so equal numbers of B cells are specific to each antigen. For all antigens in the population (*P_Ag_*), the rate (P_Ag_gN) of naive B cell (*N*) decay was set to (4.5 d)^-1^ (2a). The formation rate of naïve B cells (κ*N*) was modeled as a first-order reaction where the rate was dependent on the naïve B cell population size (5x10^7^) per antigen in the population (*P_Ag_*)) and set to (4.6x105 h)^-1^ (2b).

((3a))	$N_{j}+{Ag}_{i}\overset{\to}{\sigma_{N}\gamma_{i}Q_{ij}}B_{j}+{Ag}_{i}$

((3b))	$B_{j}+{Ag}_{i}\overset{\to}{\sigma_{B}\gamma_{i}Q_{ij}}B_{j}^{*}+{Ag}_{i}$

Naïve B cells stimulation was modeled as a second-order reaction between the antigen and naïve B cells. The rate of this reaction was determined by the base stimulation rate, immunogenicity of the antigenic site, and the binding affinity between the paratope and antigenic site. The stimulation rate was set to once per 3 days for naïve B cells. The stimulation rate for germinal center B cells was set to once per 8hrs with a maximum stimulation rate of once every 15 minutes. The rate that naïve B cells (*N*) form germinal centers (*B*) was modeled as a second-order rate equation dependent on the product of the affinity of the B cell for an antigen *i* for antigenic site *j* (Q*_ij_*), antigen (*Ag*) epitope immunogenicity (*ϒ*), and a stimulation rate multiplier *σN* (3a). The rate of germinal center B cell (*B*) stimulation was modeled as a second-order rate equation dependent on the affinity of the B cell for an antigen epitope (*Q_ij_*), antigen (*Ag*) antigenic site immunogenicity (*ϒ*), and a stimulation rate multiplier *σ_B_* (eq. 3b). Antigen (*Ag*) antigenic site immunogenicity (*ϒ*) was set to 0.8 for stalk antigenic site and 1.2 for the head antigenic sites.

((4a))	$B_{j}^{*}\overset{\to}{{rR}_{jk}}B_{j}+ B_{k}$

((4b))	$B_{j}^{*}\overset{\to}{\delta}M_{j}$

((4c))	$B_{j}^{*}\overset{\to}{\delta}P_{j}$

((4d))	$B_{j}^{*}\overset{\to}{max(\eta, gB )} 0 \eta=r (B/\kappa)$

Germinal center B cell proliferation was modeled as a first-order reaction. The product of proliferation is a single daughter B cell containing at most a single mutation from the parent genotype. The replication rate (*r*) was set to a doubling time of 8hrs. Although in reality antigen is consumed during this process, for simplicity, antigen was not consumed during B cell activation. A constant rate of differentiation (δ) for germinal center B cells was used with a probability of differentiation set to 0.1. Germinal center B cells have equal probability of differentiating into antibody secreting cells or memory B cells. Therefore, the reaction rates for differentiation (eq 4b and 4c) are depend not only on delta, but also on the probability of differentiating into either plasma or memory cell. Antibody secreting cells had a 75% chance of having a half-life of 3 days (short-lived antibody secreting cells), and a 25% chance of having a half-life of 200 days (long-lived antibody secreting cells). Affinity maturation occurs in the germinal center under high apoptotic pressure that drives the selection of higher-affinity immunoglobulin receptors. A carrying capacity for the germinal center was set to 5000 B cells. As the germinal center B cell population expands so does the rate of germinal center B cell decay. When the germinal center reaches the carrying capacity, the germinal center B cell decay rate reaches the replication rate halting further expansion of the germinal center. Germinal center B cell (*B**) proliferation was modeled as a first-order rate equation dependent on the B cell replication rate (*r*) and the probability of mutation from genotype j to genotype k (*R_jk_*) and is defined as *R_jk_* = (1 − μ)^19^(μ/3) where μ is the mutation rate (eq. 4a). A first-order rate equation was used to model memory B cells (M) (eq. 4b) antibody secreting cells (*P*) dependent on the differentiation rate of *δ* (eq. 4c). Apoptosis of germinal center B cells was modeled using a second-order equation dependent on the apoptosis rate η, which was a function of the B cell replication rate (r) and the total GC B cell population (*B*) relative to the GC carrying capacity *k* (eq. 4d).

((5a))	$P_{j}\overset{\to}{\kappa_{Ab}}P_{j}+{Ab}_{j}$

((5b))	$P_{j_{t}}\overset{\to}{{gP}_{i}}$

Antibody is produced from antibody secreting cells with a decay rate based on a half-life of once every days. Each antibody in the simulation represents a large number of real antibodies. Antibody production was dependent on presence of two types of antibody secreting cells with different decay rates. Antibody production was modeled based on the production rate *k_Ab_* (eq. 5a). The decay of short-lived and long-lived antibody secreting cell $P_{j_{s}}$ was modeled as a first-order reaction with a decay rate of *gP_i_* (eq 5b), where *i* indicates the decay rate for the antibody-secreting cell.

((6))	$M_{j}+ {Ag}_{i} \overset{\to}{\sigma_{M}\gamma_{i}Q_{ij}} B_{j}+ {Ag}_{i}$

The rate of germinal center B cell (*B_j_*) formation from Memory B cells was modeled as a second-order rate equation dependent on the affinity of the Memory B cell for an antigen epitope (*Q_ij_*), antigen (*Ag*) epitope immunogenicity (*ϒ_i_*), and a stimulation rate multiplier *σ_M_* (eq. 6). In the model, memory B cells (M_j_) give rise to germinal center B cells (B_j_) upon binding antigen (Ag_i_). Memory B cells have an increased rate of simulation compared to naïve B cells giving them a competitive advantage independent of their genotype. Additionally, given that memory B cells can arise from mutated germinal center B cells with increased affinity, memory B cells can also have greater affinity for the antigen compared to naive B cells, giving them an additional rate advantage over naïve B cells. Antigen (Ag_j_) is not consumed in this reaction. Memory B cells do not decay in the simulation.

((7a))	${Ab}_{j}+ {Ag}_{i} \overset{\to}{\rho_{i}Q_{ij}} {Ab}_{j}$

((7b))	${Ab}_{j}\overset{\to}{g_{Ab}} 0$

((8))	${Ag}_{j}\overset{\to}{g_{Ag}} 0$

Antibodies bind and remove antigen using a second-order reaction with a reaction rate that is the function of the binding affinity between the antibody paratope and antigen epitope, as well as the clearance and neutralization parameter (these values were constant between all epitopes) (eq. 7a). Intrinsic antibody (Ab) decay was based on a half-life of 10 days and modeled using a first-order rate equation dependent on the decay rate *g_Ab_* (eq. 7b). Intrinsic antigen decay was modeled based on a half-life of 12hrs and modeled using a first-order reaction dependent the antigen decay rate *g_Ag_* (eq. 8).

### Comparison of Viral Decay Kinetics

A separate exponential decay model was fit to the data for each group to assess the difference in exponential decay rate. The non-linear patterns of antigen clearance rate, with the rates appearing dependent on the current antibody levels and subsiding over time can be well depicted by the exponential decay model. This model has been used extensively in modeling viral dynamics in HIV, HBV and other diseases. The model is specified as:

$y_{ij}=(V_{1}+b_{1i})*exp\{-(\beta_{1}+b_{2i})t_{j}\}+\epsilon_{ij}$ for group SC18,

And

$y_{ij}=(V_{2}+b_{1i})*exp\{-(\beta_{2}+b_{2i})t_{j}\}+\epsilon_{ij}$ for BR07

where $y_{ij}$ is the viral load for the $i^{th}$ subject at the $j^{th}$ time point, and $t_{j}$ represents the time points in hours. Random effects, $b_{1i}$ and $b_{2i}$, are included to account for between-individual variability. The exponential decay rate is represented by $\beta_{2}$ and $\beta_{2}$ for group 1 and 2, respectively. To test for differences in the decay rates, the model was fit using PROC NLMIXED within SAS v9.4.
