## Supplemental Document for "Computer Simulations of the Humoral Immune System Reveal How Imprinting Can Affect Responses to Influenza HA Stalk with Implications for the Design of Universal Vaccines"

#### Model Sensitivity Analysis

The central finding of this report is that heterologous prime-boosting leads to boosting of highly-cross-reactive stalk specific antibodies. In order to better understand how the chosen model parameters effect stalk-specific antibody responses we performed an extensive sensitivity analysis. We first independently varied four parameters: stalk antigenic site immunogenicity, the number of antigen exposures, the number of head antigenic sites, and head antigenic site epitopic distance. These permutations provide valuable insights into the workings of the model and the humoral immune system. Additionally, we performed a Latin-hypercube-sampling-based sensitivity analysis. This analysis demonstrated the extent that each parameter effected stalk specific antibody responses.

### Individual Parameter Permutations

50 simulations were performed for each type of simulation and the count of stalk reactive antibodies was tracked. For testing the effect of the number of antigen exposures, the original model was used. For testing the immunogenicity parameter and epitopic distance parameter a two-epitope antigen was used. For testing the head antigenic site, the number of epitopes for the HA antigen was varied. The antigenic site immunogenicity parameter simulates changes in the minimum B cell receptor affinity required to stimulate a B cell. For simplicity, simulations were prime and boosted with a two-antigenic-site antigen (i.e. head and stalk) with equal epitopic distance (homologous prime-boost). As expected, as the stalk antigenic site immunogenicity was increased the stalk-specific antibodies steadily increased (SA_Fig1A). A two-fold increase in the immunogenicity parameter (0.6 to 1.2) led to a 20% increase in stalk epitope-specific antibodies on average. Increasing the number of exposures to homologous lead to an increased in stalk specific antibodies particularly during the second exposure (32% increase; SA_Fig1B). Alternatively, stalk specific antibodies decreased as the number of head epitopes increased with a 68% decrease from one to six head epitopes (SA_Fig1C). Increasing the epitopic distance led to a steady increase in stalk specific antibody (>200% increase), but plateaued when epitopic distance was increased beyond 5 (SA_Fig1D). Taken together, all parameters effected the level of stalk specific antibody, with head antigenic distance having the greatest effect.

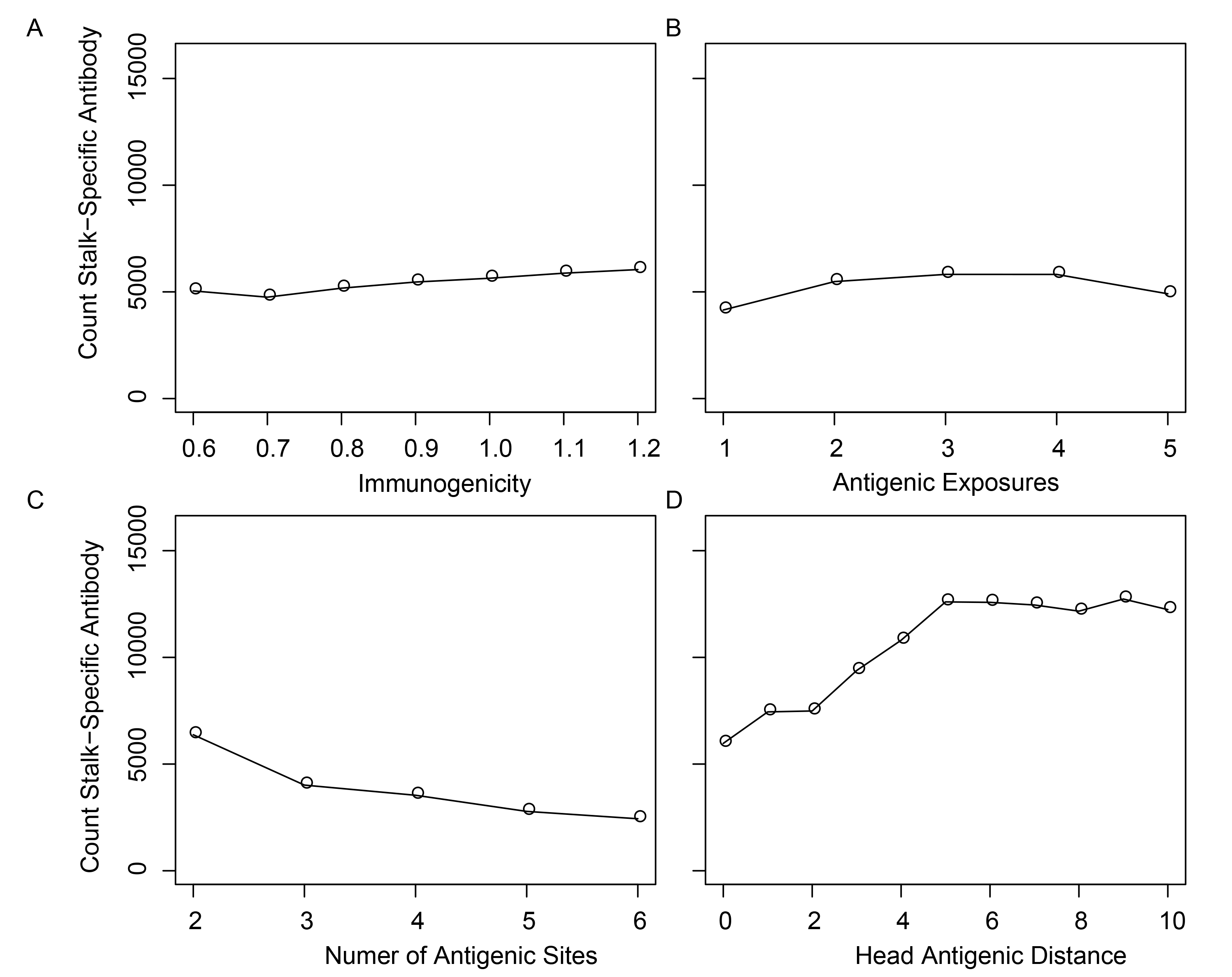

**SA_Fig1. Model Parameters on Stalk Binding Antibody Counts**

Counts of antibodies specific to stalk antigenic site were determined at day 30 post-boost of the second antigen challenge. Data is presented as the average of the 50 simulations.

(A) Stalk specific antibody counts in models with increasing stalk epitope immunogenicity parameter values. (B) Stalk specific antibody counts with different numbers of homologous antigen exposures (C) Stalk specific antibody counts with antigens that contain different numbers of head antigenic sites (D) Stalk specific antibody counts where the head epitopic distance was increased.

### **Latin Hypercube Sampling Sensitivity Analysis**

Next we assessed the extent that each parameter in the model contributes to the changes in stalk specific antibody after heterologous boosting reported here using Latin hypercube sampling. Latin hypercube sampling can be used to assess the cumulative effect of the parameters chosen for the model. We sampled values for 24 parameters used in the model. 100 Latin hypercube samples were taken from the multidimensional distribution of parameters resulting in 100 models with unique parameter values.

We found that our Latin hypercube sampling resulted in parameter values that covered multidimensional parameter space (SA_Fig2).

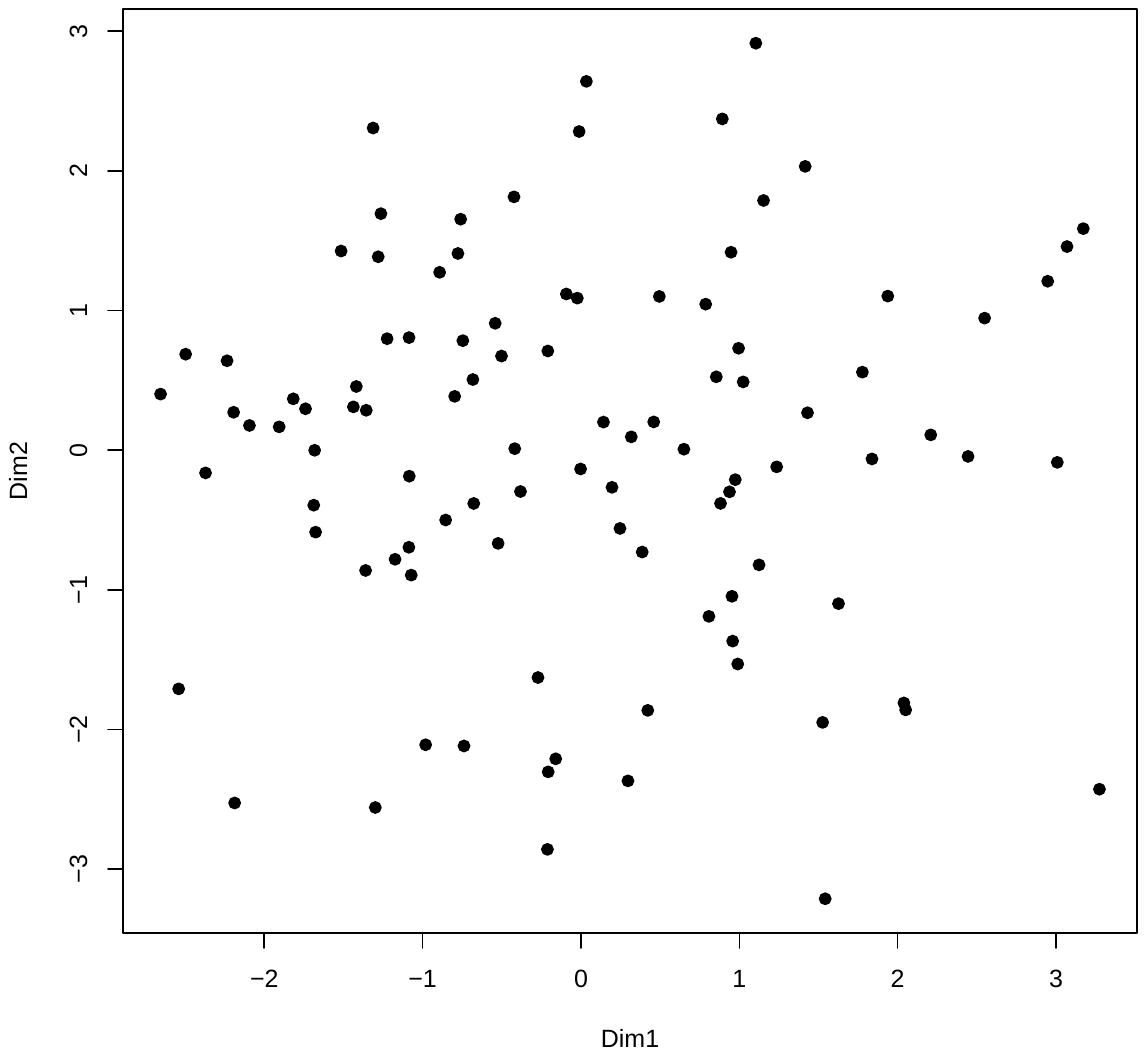

**SA_Fig2. Latin Hypercube Sampling.** Principal component analysis of the 24-dimensional parameter space derived by Latin hypercube sampling**.** Each point represents a set of 24 parameter values selected for sensitivity analysis. 90% of the variance was captured by dimensions 1 and 2.

Next, 100 unique models were created each with a unique set of parameter values. Each model was simulated 10 times and the HA stalk specific antibody levels results were recorded. A multivariate linear model (*lm* function, R base package) was used to determine the effect of each parameter value on the maximum level of stalk antibody found during the simulation.

We found that the maximum level of stalk antibody was most positively affected by the amount of initial antigen used in the model (Table 1). The Mutation Rate of the B cell receptors of germinal center B cells also positively affected the level of maximal stalk specific antibody. Ab Decay Rate, Stimulation Rate, Plasma Cell Decay Rate, and Tau also was positively correlated Stalk antibody levels, but to a lesser extent. Interestingly, although the involvement of memory B cells in the immune responds had a large impact on antibody responses (Figure 4 of main text), the B cell Enhancement Factor, which is greater in memory B cells, did not affect the maximum level of stalk antibody.

Many parameters were negatively associated with maximum stalk antibody levels where higher values significantly decreased stalk antibody levels. Antigen clearance rate, the rate in which antibodies specific to the antigen are removed from the system, was most negatively associated with stalk antibody levels. Additionally, other parameters that also influence antigen levels; Antibody production rate, Antibody Affinity, and Antigen Decay Rate; negatively influenced stalk antibody levels. Somewhat unexpectedly, the time interval between prime and boost vaccination also negatively influenced maximum antibody levels where shorter intervals lead to higher levels of stalk antibody. Interestingly, neither the B cell Carrying Capacity of germinal centers nor the Memory B Cell Differentiation Rate affected stalk antibody levels. Taken together, 14/24 (58%) of the parameters effected the maximum levels of stalk specific antibodies and many of the parameters that were directly involved in antigen amounts (e.g. initial antigen amount, antigen clearance rate, Antigen decay rate) had the greatest effect on stalk antibody levels.

**Table 1**

| ***Parameter*** | ***t-stat*** | ***p-value*** |
| --- | --- | --- |
| **Ag Initial** | **20.308** | **2.00E-16** |
| **Mutation Rate** | **18.185** | **2.00E-16** |
| **Ab Decay Rate** | **5.326** | **1.25E-07** |
| **Stimulation Rate** | **4.21** | **2.79E-05** |
| **Plasma Cell Decay Rate** | **3.578** | **0.000363** |
| **Tau** | **2.296** | **0.021893** |
| Stimulated B Cell Half Life | 1.226 | 0.220385 |
| B Cell Enhancement Factor (Memory B cells) | 0.905 | 0.365672 |
| B Cell Differentiation Rate | 0.828 | 0.408146 |
| Naïve B cell Stimulation Rate | 0.135 | 0.892702 |
| Long-Lived Plasma Cell Decay Rate | 0.014 | 0.989012 |
| **Clearance Rate** | **-8.668** | **2.00E-16** |
| **Antibody Production Rate** | **-7.043** | **3.55E-12** |
| **B cell Initial** | **-5.288** | **1.53E-07** |
| **Antibody Affinity** | **-3.937** | **8.83E-05** |
| **GC Decay Rate** | **-3.774** | **0.000171** |
| **Ag Decay Rate** | **-3.582** | **0.000357** |
| **Interval Between Vaccination** | **-3.322** | **0.000928** |
| **B cell Max Differentiation Rate** | **-3.243** | **0.001223** |
| Time From Boost Vaccination to Model End | -1.404 | 0.160587 |
| B cell Carrying Capacity | -1.399 | 0.162007 |
| B cell Affinity | -1.281 | 0.200323 |
| Basal B Cell Decay Rate | -0.916 | 0.359825 |
| Memory B Cell Differentiation Rate | -0.64 | 0.522562 |
